# Improved reference genome annotation of *Brassica rapa* by PacBio RNA sequencing

**DOI:** 10.1101/2021.11.27.470189

**Authors:** Zhicheng Zhang, Jing Guo, Xu Cai, Yufang Li, Xi Xi, Runmao Lin, Jianli Liang, Xiaowu Wang, Jian Wu

**Affiliations:** Institute of Vegetables and Flowers, Chinese Academy of Agricultural Sciences, Beijing, China

**Keywords:** PacBio-Seq, *Brassica rapa*, annotation, alternative splicing, lncRNA

## Abstract

The species *Brassica rapa* includes several important vegetable crops. The draft reference genome of *B. rapa* ssp. *pekinensis* was completed in 2011, and it has since been updated twice. The pangenome with structural variations of 18 *B. rapa* accessions was published in 2021. Although extensive genomic analysis has been conducted on *B. rapa*, a comprehensive genome annotation including gene structure, alternative splicing events, and non-coding genes is still lacking. Therefore, we used the Pacific Biosciences (PacBio) single-molecular long-read technology to improve gene models and produced the annotated genome version 3.5. In total, we obtained 753,041 full-length non-chimeric (FLNC) reads and collapsed these into 92,810 non-redundant consensus isoforms, capturing 48% of the genes annotated in the *B. rapa* reference genome annotation v3.1. Based on the isoform data, we identified 830 novel protein-coding genes that were missed in previous genome annotations, defined the UTR regions of 20,340 annotated genes and corrected 886 wrongly-spliced genes. We also identified 28,564 alternative splicing (AS) events and 1,480 long non-coding RNAs (lncRNAs). We produced a relatively complete and high-quality reference transcriptome for *B. rapa* that can facilitate further functional genomic research.

## Introduction

*Brassica rapa* is a member of the genus *Brassica*, a plant genus, widely used as vegetables (such as turnip, Chinese cabbage, pak choi, caixin), fodders and oilseed crops (e.g., turnip rape and sarson). As one of the diploid species in the genus *Brassica*, *B. rapa* was the first species for which complete genome sequencing was performed in 2011 (Wang et al., 2011). The reference genome was updated twice in 2017 and 2018 by using higher depth long reads generated from third-generation sequencing (TGS) and Hi-C technology, respectively (Cai et al., 2017; Zhang et al., 2018). In addition to the reference genome based on Chinese cabbage accession Chiifu-401-42, a number of reference genomes of yellow sarson (Belser et al., 2018) and pak choi (Li et al., 2020; Li et al., 2021) as well as a graphic genome consisting of 18 *B. rapa* accessions (Cai et al., 2021) have been published in recent years, providing valuable resources for genetic research on *B. rapa* crops. In contrast to the continuous improvement of the reference genomes of *B. rapa*, annotations for the structural integrity of genes such as the UTR regions of genes, information concerning alternative splicing (AS), alternative polyadenylation (APA), and long noncoding RNAs (lncRNAs) at the transcriptome level are still relatively scarce.

The eukaryotic transcriptome is highly complex. As two important post-transcriptional regulation mechanisms, AS and APA make significant contributions to the enrichment of the transcriptional diversity. More than 98% of human multi-exon genes occur with AS (Wang et al., 2008), and these genes tend to express many splicing isoforms simultaneously (Djebali et al., 2012). In Arabidopsis, the percentage of AS in genes is as high as 61% (Zhang et al., 2010; Marquez et al., 2012). APA can also generate transcript variants with different 3′ ends (Shen et al., 2011; Elkon et al., 2013). More than 70% of genes in Arabidopsis and 48% of expressed genes in rice contain multiple polyadenylation sites (Wu et al., 2011; Elkon et al., 2013).

Long noncoding RNAs (lncRNAs) are longer than 200 nucleotides (nt) in length, in contrast to small RNAs. LncRNAs are classified into three major types, the long intergenic noncoding RNAs (lincRNAs), intronic noncoding RNAs (incRNAs) derived from introns, and natural antisense transcripts (NATs) transcribed from the complementary DNA strand of their associated genes (Chekanova, 2015). LncRNAs have been shown to play critical regulatory roles in most eukaryotes. At present, most lncRNAs are studied in the context of protein-coding gene regulation and as a result can be functionally or mechanistically connected to mRNA expression (Wierzbicki et al., 2021). For *B. rapa*, 2,237 lncRNAs were identified from EST sequences (Paul et al., 2016), while 2,088 lncRNAs were identified based on the short-read transcriptome sequencing (Shea et al., 2019). Since StringTie assembly of transcripts was involved in these studies, mistakes could inevitably be included in the annotation of lncRNAs.

Isoform sequencing (Iso-Seq) by long reads is a revolutionary technology for plant transcriptome research. At present, Pacific Biosciences (PacBio) and Oxford Nanopore Technologies (ONT) provide full-length transcripts up to 15~25 kb for PacBio and > 30 kb for ONT from the 5′ cap to the polyA tail, thereby avoiding read reconstruction from local information (Oikonomopoulos et al., 2020). The method can retrieve most of the expressed transcripts as full-length sequences, alternative isoforms, and duplicated genes, therefore providing more accurate evidence to guide delimitation of UTRs, introns, exons, and alternative splicing junctions through genomic alignment. The continuous sequences guarantee better accuracy of gene annotation compared to expressed sequence tags (ESTs) (Adams et al., 1992), RNA-Seq (Grabherr et al., 2011), and homology inference (Jarvis et al., 2017). Because of these advantages, PacBio sequencing has been used to investigate full-length transcripts and to construct reference gene sets in plant species including rice (Zhang et al., 2019), maize (Li et al., 2014b), moso bamboo (Wang et al., 2017), and *Brassica napus* (Yao et al., 2020).

The present study analyzed the full-length transcriptome of the Chinese cabbage accession Chiifu-401-42, the strain used for previous *B. rapa* reference genome assembly. The genome was derived from five different tissues using PacBio SMRT sequencing. The updated genome annotation provides a reference gene set for future functional genomic studies. Furthermore, improved gene annotations could contribute to better understanding of the complexity of the *B. rapa* genome and serve as a reference sequence for gene functional studies.

## Materials and Methods

### Plant materials

Chinese cabbage accession Chiifu-401-42 (referred as Chiifu), which had been used for *B. rapa* reference genome assembly, was used in this study. Total RNA was isolated from 100 germinated seeds. The germinated seeds were sown in pots filled with peat and vermiculite at a ratio of 2:1. The seedlings were grown in a greenhouse located in the Haidian district in Beijing. After 21 days of growth, young leaves were harvested from single seedlings for RNA isolation. Then, the seedlings were transferred to a growth chamber for vernalization treatment (12 h light/12 h dark, 4°C day/night temperature). Young leaves were collected from single seedlings at 14/28/35/50 days after the plants were placed in the vernalization treatment. Total RNA was isolated from each sample and mixed in equal amounts for library construction (Library I).

In addition, germinated seeds were sown in pots filled with peat and vermiculite at a ratio of 2:1 and seedlings were grown in the same greenhouse as mentioned above. The shoots for single plants were harvested at 21 days after sowing (LNV). Then the remaining seedlings were transferred into cold treatment and the shoots were collected at the time of 7/14/21/28 days after transferring into a cold condition (LV). Plants at the stage of heading formation were sampled for leaves and petioles as well as roots. In addition, the inflorescence was harvested from flowering plants. Total RNA was isolated separately for LNV, LV, roots, heading leaves, petioles, and the inflorescence. Equal amounts of each RNA sample were mixed for library construction (Library II).

### Library preparation and Iso-Seq

The Iso-Seq libraries were prepared according to the Isoform Sequencing protocol (Iso-Seq) using the Clontech SMARTer PCR cDNA Synthesis Kit and the BluePippin Size Selection System protocol as described by Pacific Biosciences (PN 100-092-800-03). Library I and Library II were each constructed with random-size cDNA fragments > 4 kb in length. The libraries were subsequently sequenced on the PacBio RS II platform using P6C4 polymerase enzyme.

### RNA-seq library preparation and sequencing

Total RNA isolated from leaves was sent to Novogene (www.novogene.com) for transcriptome sequencing. Briefly, mRNA was purified from total RNA using poly-T oligo-attached magnetic beads. Fragmentation was carried out using divalent cations under elevated temperature in First Strand Synthesis Reaction Buffer (5X). First strand cDNA was synthesized using random hexamer primers and M-MuLV Reverse Transcriptase, and then use RnaseH was used to degrade the RNA. Second strand cDNA synthesis was subsequently performed using DNA Polymerase I and dNTP. The remaining overhangs were converted into blunt ends via exonuclease/polymerase activities. After adenylation of 3′ ends of DNA fragments, adaptors with hairpin loop structure were ligated to prepare for hybridization. In order to preferentially select cDNA fragments 370~420 bp in length, the library fragments were purified with the AMPure XP system (Beckman Coulter, Beverly, USA). Then PCR was performed with Phusion High-Fidelity DNA polymerase, Universal PCR primers and Index Primer. Finally, PCR products were purified (AMPure XP system) and library quality was assessed on an Agilent Bioanalyzer 2100 system. Five qualified libraries (NV, V1, V2, V3, and V4) were sequenced on an Illumina NovaSeq platform and 150 bp paired-end reads were generated.

### Data collected from public databases

The *B. rapa* reference genomic sequences v3.0 (Zhang et al., 2018) and genome annotation (v3.1) (http://39.100.233.196:82/download_genome/Brassica_Genome_data/Brara_Chiifu_V3.1/) were downloaded from BRAD (Chen et al., 2021). The genomic sequences and protein sequences of three related species—*A. thaliana* (TAIR10), *B. oleracea* (JZS_v2.0), and *B. nigra* (NI100_v2.0) —were also downloaded from BRAD (http://brassicadb.cn/). In addition, 31G published Illumina-based RNA-seq datasets of Chiifu were collected form NCBI (https://pubmed.ncbi.nlm.nih.gov/), including data generated from a variety of tissues (Supplementary Table 1).

### PacBio sequencing data processing and error correction

The PacBio-seq raw reads were filtered into circular consensus sequence (CCS) subreads using SMRTLink (https://github.com/PacificBiosciences/IsoSeq/). After quality evaluation, CCS subreads were processed to generate the ROIs with a minimum full pass of 1 and minimum accuracy of 95. The pipeline then classified the ROIs in terms of full-length non-chimeric and non-full-length reads. This was done by identifying the 5′ and 3′ adapters used in the library preparation as well as the poly(A) tail. Only reads that contained all three in the expected arrangement and did not contain any additional copies of the adapter sequence within the DNA fragment were classified as full-length non-chimeric (FLNC) reads. Then, the FLNC reads were corrected using LoRDEC (version 0.9) (Salmela and Rivals, 2014) and an RNA -seq dataset consisting of 171,308,229 pairs of reads form six different tissues (callus, root, stem, leaf, flower and silique) (Tong et al., 2013).

### Alignment of PacBio reads to the *B. rapa* reference genome and collapse

All FLNC reads were mapped to the *B. rapa* reference genome using minimap2 (version 2.15) (Li, 2018) software with the options: -ax splice, -uf, -C5, -O6, and 24 - B4. Only the best alignments (the largest alignment with the lowest number of mismatches against the reference genome) were kept. Then the retained FLNC reads were collapsed into full-length non-redundant consensus isoforms using a Python script (collapse_isoforms_by_sam.py) in the Cupcake program (version 19.0.0) (https://github.com/Magdoll/cDNA_Cupcake/wiki/Cupcake:-supporting-scripts-for-Iso-Seq-after-clustering-step#collapse) with parameters of coverage > 0.90 and identity > 0.90.

### Expression level estimation

We used fastp (Chen et al., 2018) software with default options of quality control of the RNA-seq data. The high-quality clean reads were aligned to the *B. rapa* reference genome using HISAT2 (version 2.1.0) (Pertea et al., 2016). First, we built a HISAT2 index and aligned the reads for each sample to the reference genome with the default options. Then, we sorted and converted the SAM files to BAM files using SAMtools (version 1.6.3) (Li et al., 2009). Finally, the quantification of isoforms and genes was performed using StringTie (version 2.0.4) (Pertea et al., 2015) based on the GTF/GFF3 annotation file generated by PacBio sequencing, a gene annotation file, and a BAM file containing aligned reads information. The expression levels were measured and normalized as transcripts per kilobase of exon model per million mapped reads (TPM).

### Novel gene annotation

The isoforms that had no overlap with any annotated regions were defined as candidate novel transcripts. Then, their coding ability was predicted using TransDecoder (version 5.5.0). We selected the longest read as the representative sequence for each isoform to search the following databases. For the tblastx analysis, we built a database consisting of all plant cDNAs of protein-coding genes from Ensembl Plants, release 51 (ftp://ftp.ensemblgenomes.org/pub/plants/release-51/, last accessed on 20210505). For blastx, we used the Swiss-Prot database, the current release (ftp://ftp.uniprot.org/pub/databases/uniprot/current_release/knowledgebase/complete/, last accessed on 16 Jun 2021) containing 565,254 protein sequences. Finally, we ran tblastx and blastx with the criterion of e-value ≤ 10e-6.

### Identification of alternative splicing events

The alternative splicing (AS) events of *B. rapa* were identified by employing the Astalavista (version 4.0) (Foissac and Sammeth, 2015) algorithm with default options based on the assembled GTF file from the Iso-Seq. This study focused on four major types of AS events that were classified by the codes: IR (0, 1^2-), AA (1-, 2-), AD (1^, 2^) and ES (0, 1-2^) (Foissac and Sammeth, 2015). The statistical analysis was performed using custom Perl scripts.

### Identification of lncRNAs from PacBio data

Putative protein-coding RNAs were filtered using the min-length of sequence and min-length of the ORF thresholds. Non-protein-coding RNA transcripts longer than 200 bp and ORFs with length shorter than 100 bp were sorted as lncRNA candidates using four computational approaches, namely CPAT (Wang et al., 2013) (Coding Potential Assessment Tool), CPC (Kong et al., 2007) (Coding Potential Calculator), PLEK (Li et al., 2014a) (predictor of long non-coding RNAs and messenger RNAs based on an improved k-mer scheme), and LGC (Wang et al., 2019a), all of which could distinguish protein-coding and non-coding transcripts. Only reads having consistent results from at least two software programs were considered as lncRNA candidates. To further verify the credibility of lncRNA, the remaining reads were searched against the Swiss-Prot database using blastx (*E*-value, 10e^−6^).

### Analysis of differential expression of lncRNAs (DELs)

The HISAT2 soft was used to align RNA-seq data against the *B. rapa* reference genome. The output SAM file was then sorted and converted to a BAM file using SAMtools. Next, the htseq-count script of HTSeq (Anders et al., 2015) (version 0.9.1) was used to calculate the number of reads mapping to each lncRNA, with the opitions -f bam and -s no, and the output count file was used for dowmstream analysis. The DESeq2 package (Love et al., 2014) (version 1.34.0) was used to identify the DELs between NV and V samples with a *P*-value < 0.05 and |fold change| > 2.

### Validation by RT-PCR

Total RNA was extracted using TRIzol reagent (TransGen) as described in the user’s manual. For cDNA synthesis, 2 μg of total RNA was treated with gDNA Remove (TransGen) and then used to synthesize the first-strand cDNA using TransScript One-Step gDNA Removal and cDNA Synthesis SuperMix (TransGen). For PCR validation of AS events, split genes and the non-coding exons of *BrSOC1*.*1* and *BrSOC1*.*2*, 3 μl of the cDNA was used in a reaction volume of 50 μl using 2 x Rapid Taq Master Mix (Vazyme). The specific primers were designed by the Primer-BLAST Tools of NCBI (https://www.ncbi.nlm.nih.gov/tools/primer-blast/). PCR conditions were 3 min at 95 °C followed by 40 cycles at 95 °C for 15 s, 55 °C for 15 s and 72 °C for 15 s, and final extension at 72 °C for 5 min. The PCR amplification was visualized on 2% agarose gel.

### Data availability statement

The dataset generated has been deposited into the brassica database (http://brassicadb.cn/), to allow easy access to the data and to facilitate for the exploration of the annotation and full-length isoforms using a genome browser.

## Results

### FL sequencing of *B. rapa* and isoform detection

In this study, we combined equal amounts of total RNA from five different tissues to build two cDNA libraries of random sized fractionated (see Methods) for acquiring as many full-length transcripts as possible by single-molecule long-read sequencing. In total, 1,323,470 reads of inserts (ROIs) were generated and filtered using the SMRT Link software (https://github.com/PacificBiosciences/IsoSeq/), among which 778,561 were identified as full-length non-chimeric (FLNC) reads based on the presence of both 5′ and 3′ signals plus the polyA tails.

As the mapping accuracy is affected by read quality, especially for the correct identification of splicing junctions, we performed read error correction using short-read Illumina data from the SRA database (Supplementary Table 1) with LoRDEC (Long-Read DBG Error Correction, v0.9) (Salmela and Rivals, 2014). The corrected reads had a higher Benchmarking Universal Single-Copy Orthologs (BUSCO) (Seppey et al., 2019) completeness value (97.2%) compared to the uncorrected reads (88.7%) (Supplementary Figure 1).

Next, the corrected FLNC reads were mapped to the *B. rapa* genome using minimap2 (version 2.15) (Li, 2018). In total, 753,041 corrected reads were uniquely mapped to reference genome, including 17,962 reads being mapped to genomes of chloroplasts and mitochondria that were excluded from further analysis. Following previous studies, we defined collapsed reads as isoforms, and read clusters as loci (Wang et al., 2019b). Through this, the alignments were collapsed into 92,810 full-length non-redundant consensus isoforms belonging to 26,163 loci. The mean length of isoforms was 2,171 bp, longer than that of annotated genes (1,155 bp).

In addition, we observed that of the 35.5% (32,962) of full-length non-redundant consensus isoforms with less than 90% coverage of annotated genes, 3,683 were located in intergenic regions. The isoforms that could not be mapped to any annotated genes might be novel genes that had not been identified in previous studies. The low-coverage isoforms may be ncRNAs or different transcripts due to variable splicing, or possibly site mutations. For this reason, the Iso-Seq data in this study better reflected the actual transcription of genes; compared to previous annotations, this analysis provides deeper information concerning RNA processing.

### Improving *B. rapa* genome annotation by PacBio sequencing

We compared the 92,810 non-redundant isoforms against the gene set from the previous genome annotation. The results showed that 59,848 isoforms (64.49%) could match (coverage > 90%) to 22,830 annotated genes, including 58,180 isoforms uniquely overlapped to 21,681 annotated genes (Figure 1A, left). Among these 21,681 genes, 8,980 were covered by one isoform, 5,104 genes were covered by two isoforms, 2,827 genes by three isoforms, 1,596 genes by four isoforms, and 3,174 genes by more than four isoforms (Figure 1A, right). For the 58,180 uniquely mapped isoforms, the coding region sequences and corresponding amino acid sequences were predicted by TransDecoder software. <u>Using</u> the above methods, the UTR regions of 20,340 annotated genes were defined, the remaining 1,341 annotated genes were not defined because their isoforms were considered lacking both initiation codons and termination codons by prediction software.

**Figure 1.**
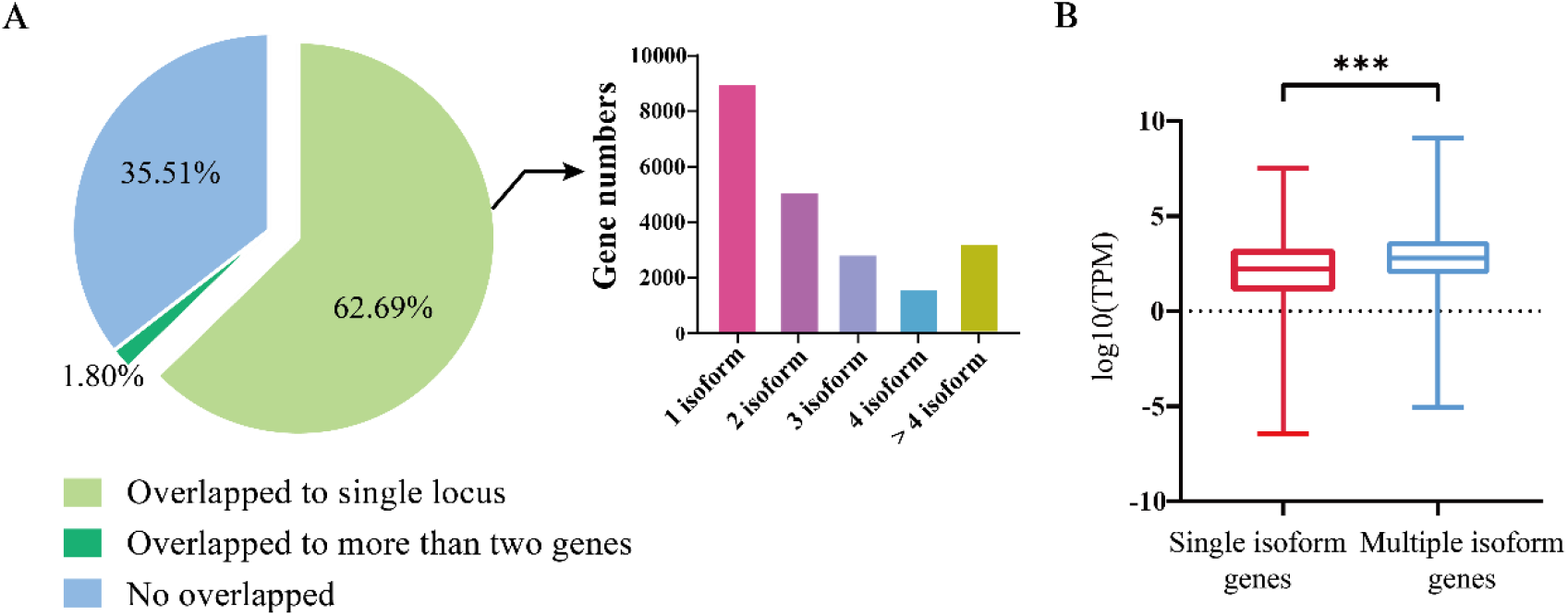
Identification and characterization of isoforms and the genes involved. **(A)** Left, division of the FLNC isoforms based on their genome mapping status. The percentages of isoforms in each group are depicted in the pie chart. Right, the distribution of genes covered by different number of isoforms. **(B)** The expression levels of genes covered by single isoforms and multiple isoforms.

We then compared the expression levels between the genes covered by multiple isoforms and those covered by a single isoform. A total of 31 GB of transcriptome data was downloaded from the NCBI SRA Database for comparison (Supplementary Table 1). It turned out that the genes covered by multiple isoforms were expressed at higher levels than the genes covered by a single isoform (*t*-test, *P*-value < 2e^−16^, Figure 1B).

The *B. rapa* reference genome was annotated at the coding level in previous annotation version; therefore, non-coding exons were neglected. By aligning isoforms to the genome annotation, we detected non-coding exons for 7,436 annotated genes and corrected their gene model in v3.5. For example, there are two transcriptional forms of the *SOC1* gene (*AT2G45660*) in *A. thaliana* (Araport11, (Cheng et al., 2017) (Figure 2A), the main form (*AT2G45660.1*) is 3,620 bp in length and consists of eight exons, and the other form (*AT2G45660.2*) is 3,547 bp in length with seven exons caused by alternative splicing between the original sixth and seventh exons (Figure 2A). The first exon is non-coding and translation starts from the second exon to the eighth exon of *AT2G45660.1*. *B. rapa* contains three copies of *SOC1, BrSOC1.1* (*BraA05g005300.3.1C*), *BrSOC1.2* (*BraA04g032590.3.1C*), and *BrSOC1.3* (*BraA03g023940.3.1C*). Transcripts of two copies, *BrSOC1.1* and *BrSOC1.2*, were captured by Iso-Seq. The alignment results showed the previously annotated *BrSOC1.1* containing a non-existing exon 7 while lacking the first exon compared with the full-length transcripts by Iso-Seq (Figure 2B). The annotated *BrSOC1.2* gene also lacked of the first exon (Figure 2C). ORF prediction showed that the CDS starts from the second exon of both *BrSOC1.1* and *BrSOC1.2*, in line with that of *A. thaliana*. We then designed primers to amplify fragments across the first and second exon for *BrSOC1.1* and *BrSOC1.2*. The results of reverse transcription (RT)-PCR confirmed the presence of the first exon (Figure 2 B, C). In addition to the two fragments in 277 bp and 409 bp in length in the amplification of *BrSOC1.2*, a fragment approximately 300 bp in length was also amplified, indicated that there could be an extra transcript model of this gene.

**Figure 2.**
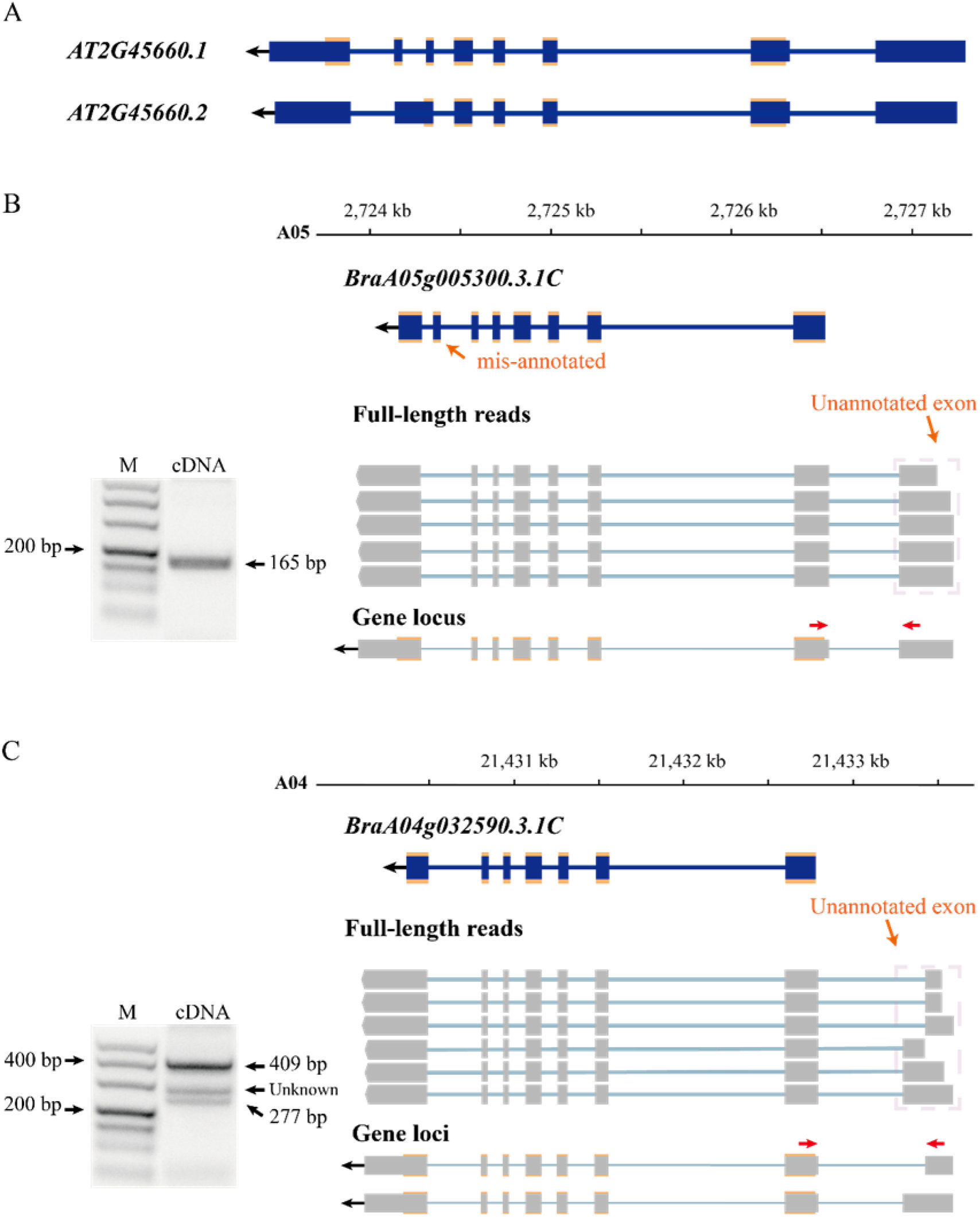
Improving gene annotation by Iso-seq data. **(A)** The gene structure of *SOC1* in *Arabidopsis thaliana* (Araport11). The blue boxes indicate exon and the croci boxes indicate the CDS of *AtSOC1*. **(B)** and **(C)** RT-PCR validation of *BrSOC1.1* (*BraA05g005290.3.5C*) and *BrSOC1.2* (*BraA04g032780.3.5C*) structure. Forward and reverse primers are shown as red arrows. The schematic representation of genes and full-length reads is shown. The gene structure of the previously annotated *BrSOC1.1* and *BrSOC1.2* (blue) and the transcripts detected by Iso-Seq (gray). The gene locus shows *BrSOC1.1* and *BrSOC1.2* structures in v3.5; the blue and gray boxes indicate exons and the croci boxes indicate CDS. The PCR products of unknown length may indicate that the type was not captured.

By aligning isoforms to the reference gene annotation, we detected the isoforms covering adjacent annotated genes in previous annotations, suggesting that a single gene was mis-annotated as multiple genes (Figure 3A). In total, 1,540 isoforms simultaneously were found overlapping two to four annotated genes. We then filtered these isoforms according to four criteria, i) containing intron in length longer than 10 kb; ii) poor alignment quality (MAPQ < 10); iii) compared with the overlapped annotated genes, the first or last intron of the isoform being longer than the length of any intron of the annotated gene; iv) the length of the first or last exon being less than 5% of the total length of the isoform. The isoforms that satisfied any one of these four criteria were removed. After filtering, 1,175 isoforms were defined as candidate sequences. Of these, 1,052 isoforms were predicted to be protein-coding, indicating that these 1,052 isoforms had been mis-annotated in the reference gene annotation. Further transcriptional analysis of RNA-seq data using second-generation sequencing showed that 1,052 isoforms reached a level above 1 TPM, indicating that these isoforms were true transcripts. In order to further validate these gene models, we randomly selected three isoforms to design primers (Supplementary Table 2) that could amplify fragments spanning wrongly-split gene pairs and performed RT-PCR using RNA isolated from Chiifu leaves (Figure 3A). The successful amplification of fragments further confirmed that these genes were wrongly split. Using the above analysis, we corrected 886 wrongly-split genes in the previous gene annotation into 427 genes in v3.5.

**Figure 3.**
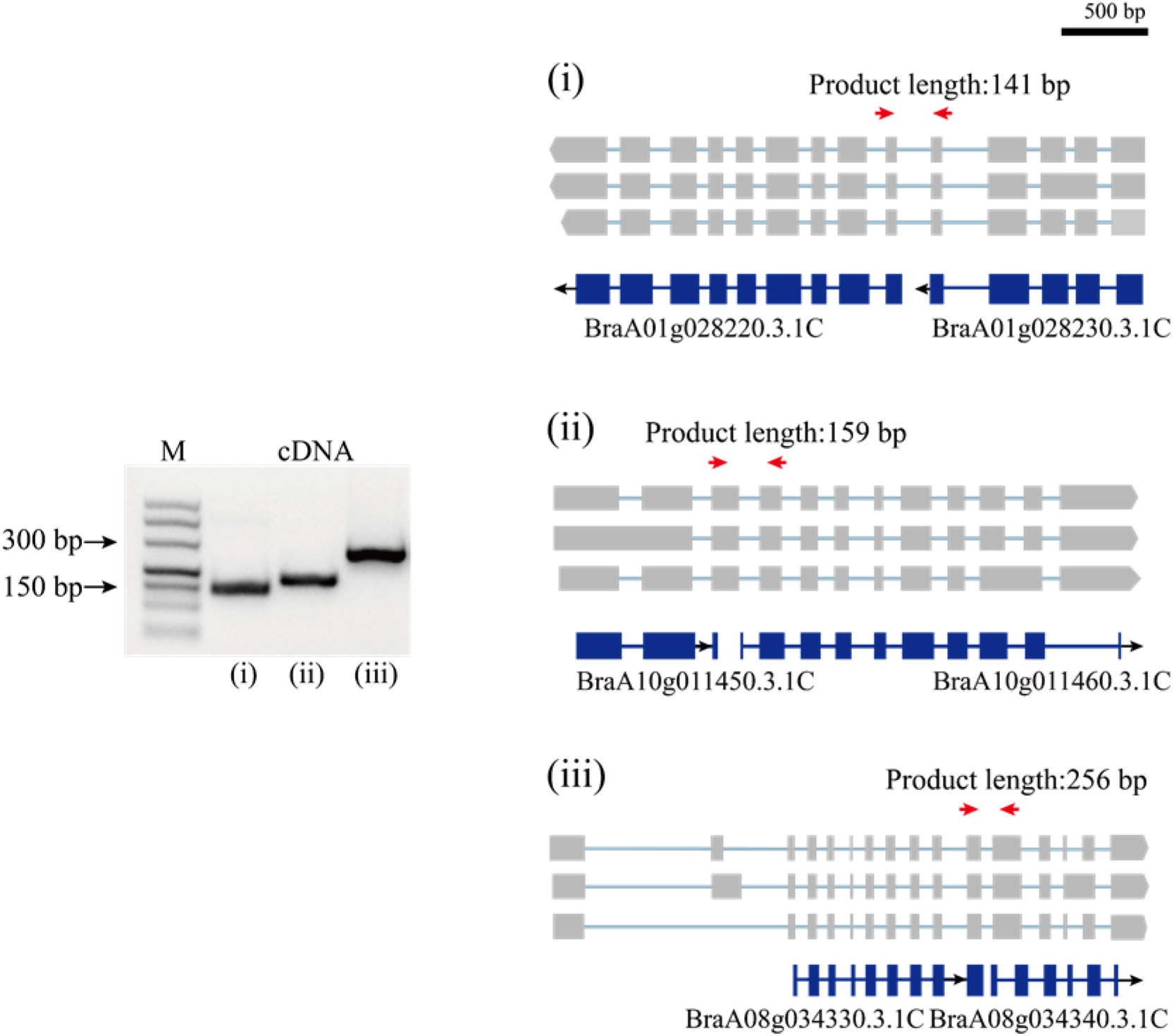
PCR validation of wrongly split genes. RT-PCR validation of wrongly split genes. The schematic representation of genes and full-length reads is shown. Gene structures in blue shows mis-annotated genes in the previous annotation. The gray boxes indicate isoform exons. Forward and reverse primers are shown as red arrows.

Among the 92,810 non-redundant isoforms, 3,683 isoforms were generated from 1,707 novel loci, that were absent from the reference genome annotation. Of these, 1,829 isoforms were predicted to have coding capability by TransDecoder software and were defined as candidate novel protein-coding transcripts. To further confirm these novel loci, we searched the presence of these novel isoforms in other organisms using blast (v2.9.0). Among the 1,829 isoforms, 1,825 isoforms were significantly matched against a collection of plant protein-coding gene cDNAs by tblastx (*E-*value < 10e^−6^), while 1,468 isoforms were detected in the blastx search against Swiss-Prot proteins (*E-*value < 10e^−6^). Notably, all 1,468 isoforms detected by blastx completely overlapped with the 1,825 isoforms detected by the tblastx search (Figure 4A). Given this high ratio of overlap, the 1,825 isoforms from 830 loci detected in the PacBio sequencing data likely represented transcripts from novel protein-coding genes. Through the above analysis, 830 novel protein-coding genes were supplemented to *B. rapa* genome annotation v3.5.

**Figure 4.**
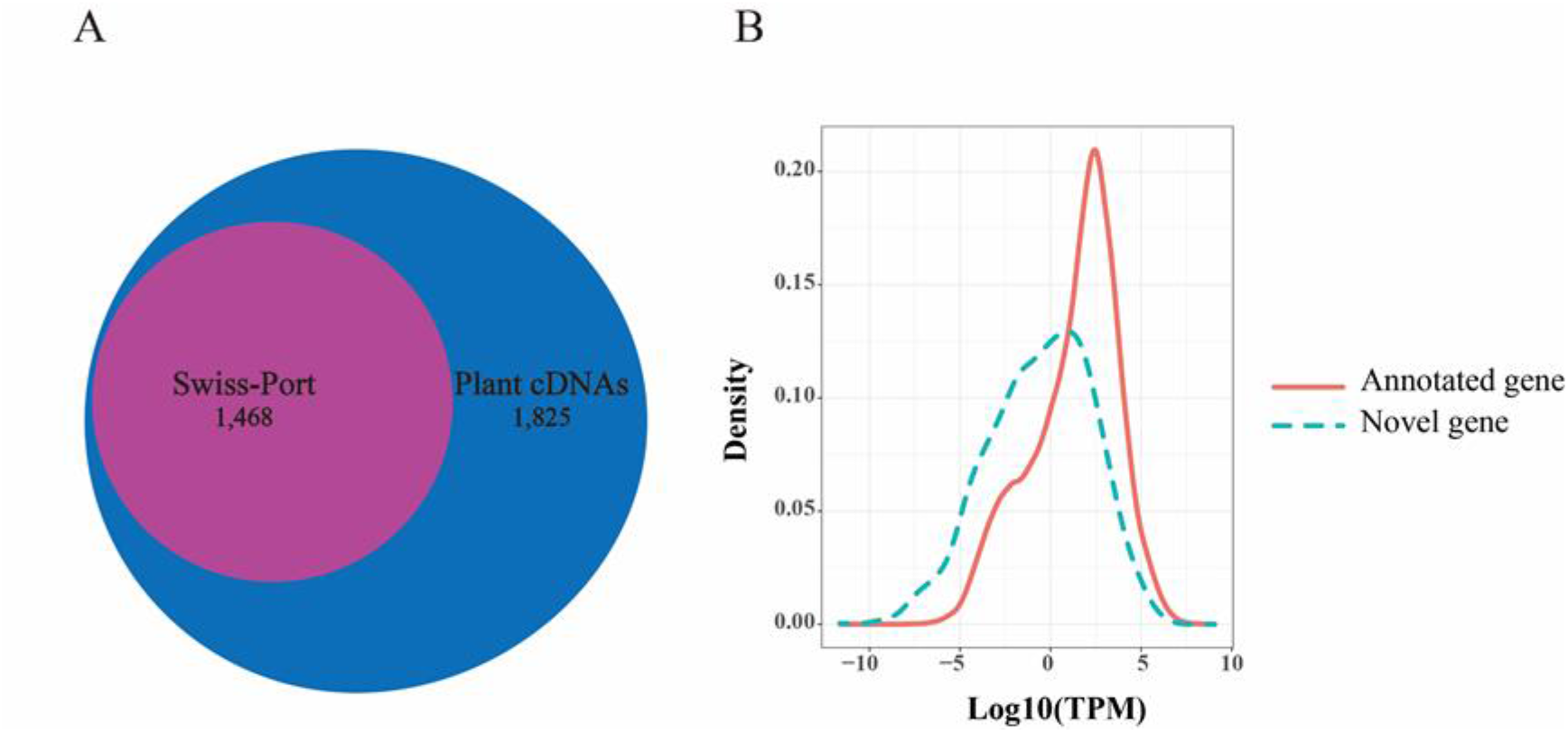
Novel genes identified in Iso-Seq data. **(A)** The number of novel genes in *B. rapa* that showed significant sequence similarity in a blastx search against Swiss-Prot proteins or in a tblastx search against plants cDNAs. **(B)** Distributions of transcripts per million (TPM) values of novel genes (green dotted line) and annotated genes (red line).

In addition, we found that the average expression levels of these novel genes (1,825 isoforms, TPM = 7.322) were lower than those of the previously annotated genes (46,878 gene, TPM = 21.048) based on Illumina short reads (*t*-test, *P*-value = 7.60e^−67^, Figure 4B). This suggested that a low expression level might be one of the reasons for missing these genes in the previous annotation. In contrast, the average exon number of novel genes (mean = 7.43) was significantly greater than that of the previously annotated genes (mean = 5.13) (*t*-test, *P*-value = 1.045e^−05^). This indicated that the number of exons may also affect gene annotation.

### Analysis of alternative splicing events and splice isoforms

Alternative splicing (AS) is prevalent in most plant genomes. One of the most important advantages of PacBio sequencing is its ability to identify AS events by directly comparing different isoforms of the same individual gene. From the 753,041 FLNC reads collapsed to 92,810 non-redundant consensus isoforms, we identified 28,564 AS events using the Astalavista program (Foissac and Sammeth, 2015) without transcriptome assembly, in order to avoid possible artificial results.

The 28,564 AS events were then clustered into four distinct types: alternative 5′-donor (AD, 3,739), alternative 3′-acceptor (AA, 7,098), exon skipping (ES, 1,310) and intron retention (IR, 16,417) (Figure 5A). Consistent with previous studies, intron retention occupied the majority of AS events (Campbell et al., 2006). However, since there are no ribosome occupancy data available, we cannot exclude the possibility that some of the retained introns were found in incompletely processed nuclear transcripts. Among the 21,681 annotated genes by Iso-Seq, 8,980 possessed a single transcript, while 12,701 possessed two or more transcripts, producing 59,596 isoforms in total, with an average of 4.69 isoforms per gene. The genes with multiple isoforms had more exons than single isoform covered genes (*t*-test, *P*-value < 2e^−16^) (Figure 5C). Only 28.9 % of genes with a single transcript contained more than five exons, however, 53.0 % of genes with alternative splicing had more than five exons (Supplementary Table 3). There were cases where the same AS type occurred at different sites within a gene locus, generating different isoforms. Multiple AS events most frequently happened for IR (Figure 5B).

**Figure 5.**
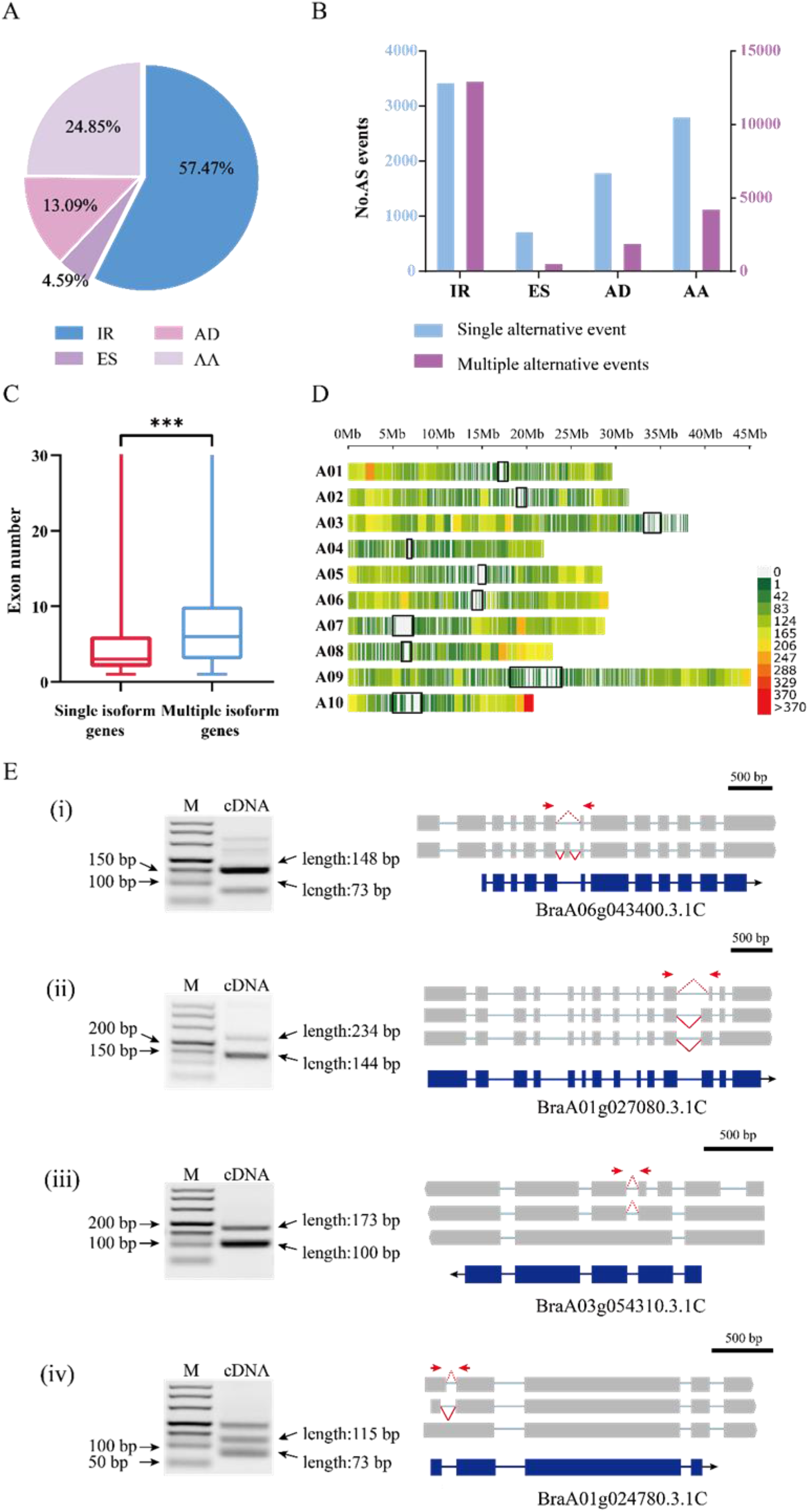
Analysis of alternative splicing events and splice isoforms. **(A)** Proportions of four AS types classified according to biogenesis. IR: intron retention; AD: alternative 5′-donor; ES: exon skipping; AA: alternative 3′-acceptor. **(B)** In the histograms, transcript variants were sorted into isoforms by single alternative event (blue) and multiple events (red). **(C)** The exon numbers of genes covered by multiple isoforms and single isoforms. **(D)** The density distribution of alternative splicing on chromosomes. Gene density was calculated in a 1 Mb sliding window. The black blocks indicate the centromere sites. **(E)** The RT-PCR validation of AS genes for four typical types. (i) exon skipping; (ii) alternative 3′-acceptor; (iii) intron retention; (iv) alternative 5′-donor. Forward and reverse primers are shown as red arrows. The blue boxes indicate gene CDS, and the gray boxes indicate isoform exons. The red dashed and solid lines indicate different splicing events.

In addition, we investigated the distribution of AS in the three subgenomes of *B. rapa*. The results showed that there was no significant difference between the frequency of AS on the dominant subgenome LF (29.8%) and that on the two submissive subgenomes MF1 (25.4%) and MF2 (27.6%), especially in cases where the gene number was higher on the LF subgenome. However, when considering only three-copy genes, we found that AS occurred significantly more on LF (228) than on MF1 (162) or MF2 (165) (ANOVA, *P*-value = 0.026). For the two-copy genes, AS frequency was not significantly different between LF and MFs or between the two MFs (Supplementary Table 4).

We set a sliding window of 1 Mb to investigate the distribution of variable splicing events on chromosomes. The results showed that the variable splicing events mostly occurred in the regions far from the centromere sites (Figure 5D). On one arm of chromosome A10, the end region showed a very high frequency of AS; in contrast, on the short arm of chromosome A03, AS occurred very rarely. Chromosome A03 is acrocentric, with a heterochromatic upper (short) arm bearing the nucleolar organizer region (NOR) (Mun et al., 2010).

To verify the AS events detected above, we randomly selected four genes and designed primers (Supplementary Table 2) for the AS regions and performed RT-PCR using RNA isolated from Chiifu leaves. The amplified fragments were consistent in length with the splicing isoforms identified from Iso-Seq data (Figure 5E).

### LncRNA identification in Iso-Seq data and differentially expressed lncRNAs after vernalization

LncRNAs, a class of non-coding RNAs are essential regulators of a wide range of biological processes. In this study, we used four computational approaches, namely CPAT (Wang et al., 2013), CPC (Kong et al., 2007), PLEK (Li et al., 2014a), and LGC (Wang et al., 2019a), to identify isoforms without coding capability. We eliminated the isoforms with ORFs exceeding 100 amino acids or sequences shorter than 200 bp. To obtain a high-confidence set of lncRNAs, we kept the predicted lncRNA only if it was predicted as non-coding by any two of the above four programs. The analysis generated a total of 11,054 candidate lncRNAs. Then, we screened their homology with Swiss-Prot proteins using blastx (*E-*value < 10e^−6^) and eliminated 7,655 isoforms homologous to proteins. Among the remaining 3,399 isoforms, 1,919 isoforms could not be defined or classified because their sequences did not satisfy the definition of known lncRNAs. Finally the retaining 1,480 lncRNAs (Supplementary Table 5) were clustered into three main types according to their distribution on chromosomes: 1,148 isoforms mapped to intergenic regions of the genome were defined as long intergenic noncoding RNAs (lincRNAs); 289 isoforms mapped to the anti-sense strand of a known gene locus were defined as natural antisense transcripts (NATs); 43 isoforms mapped to the intronic regions of the genome were defined as intronic noncoding RNA (incRNAs) (Figure 6A). The average length of the lncRNAs was 1,402 bp (range, 0.2-7.8 kb), which was longer than the 497 bp from a previous study (Paul et al., 2016). The length of most lncRNAs was between 200 and 2,000 bp (82.5%), and the number of lncRNAs decreased as the length increased (Supplementary Figure 2a). Most of the lncRNAs (1,201) consisted of one or two exons (Supplementary Figure 2b).

**Figure 6.**
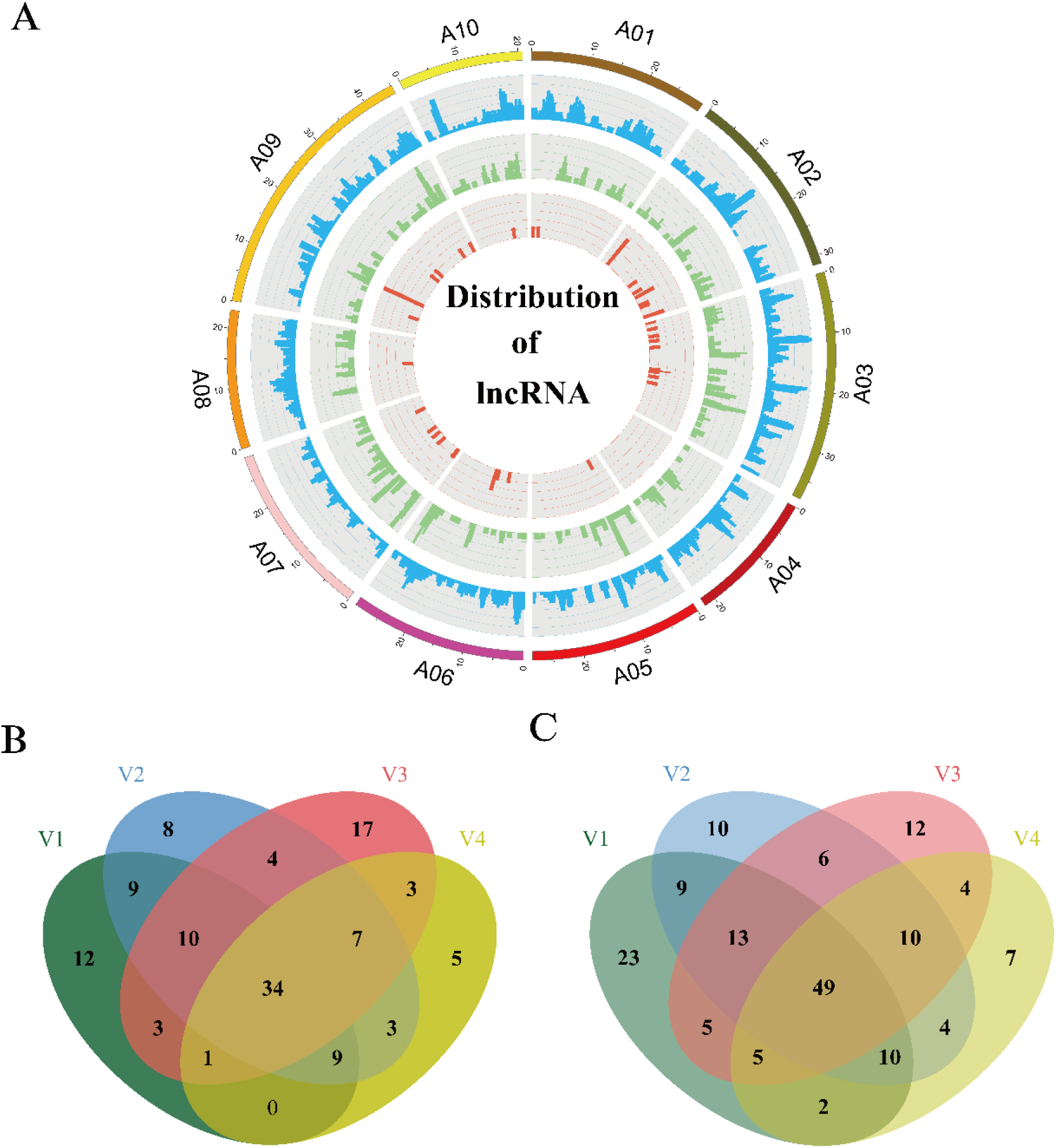
Distribution of lncRNAs and differently expression analysis. **(A)** Distribution of lncRNAs along each chromosome. From inside to outside, red, green, and blue represents intronic noncoding RNAs (incRNAs), natural antisense transcripts (NATs), and long intergenic noncoding RNAs (lincRNAs). The abundance of lincRNAs, NATs, incRNAs in physical bins of 1 Mb for each chromosome (generated using Circos). **(B)** and **(C)** Venn diagram respectively depicts common and unique lncRNAs up-regulated (B) or down-regulated (C) under different vernalization conditions. V1, V2, V3, V4 indicating the plants vernalized for 7, 14, 21, 28 days, respectively.

LncRNAs have specific functions in plant development and various biological processes, and they are reported to be involved in the response to environmental stimuli (Rinn and Chang, 2012). For example, vernalization in *Arabidopsis* influenced by lncRNAs *COOLAIR* (an antisense lncRNA) and *COLDAIR* (an incRNA) has been characterized (Swiezewski et al., 2009; Heo and Sung, 2011). Therefore, we investigated the expression profiles of lncRNAs during the vernalization process. We performed RNA-seq for the aboveground parts of Chiifu plants vernalized for 7, 14, 21 and 28 days (V1, 2, 3, 4) and without treatment (NV). We used the DESeq2 (v1.26.0) package to compare the expression levels of lncRNAs between vernalized plants (V) and non-vernalized plants (NV). A total of 470 lncRNAs were detected as being differentially expressed under the criteria of *P*-value < 0.05 and |fold change| > 2 during vernalization. Of these, 78 (V1), 84 (V2), 79 (V3), and 62 (V4) lncRNAs were up-regulated (Figure 6B), and 116 (V1), 111 (V2), 104 (V3), and 91 (V4) lncRNAs were down-regulated (Figure 6C). A total of 83 differentially expressed lncRNAs continuously changed throughout the 28 days of vernalization, 34 being up-regulated and 49 being down-regulated (Figure 6B, C). These 83 lncRNAs comprised 20 NATs (10 up-regulated and 10 down-regulated), three incRNAs (all up-regulated), and 60 lincRNAs (21 up-regulated and 39 down-regulated).

Three cold-responsive noncoding RNAs (COOLAIR (Swiezewski et al., 2009), COLDAIR (Heo and Sung, 2011), and COLDWRAP (Kim and Sung, 2017)) were reported within the *FLC* locus in *A. thaliana* (Figure 7A). Five NATs (Li et al., 2016) and a single NAT (Shea et al., 2019) were detected at the *BrFLC2* locus of *B. rapa* (Li et al., 2016; Shea et al., 2019) (Figure 7B) in two independent studies. We examined the noncoding RNAs for the four *BrFLC* loci, and only detected one NAT of length of 1,022-bp (*PB25365.2*) on the *BrFLC1* (*BraA10g027780.3.1C*) locus (Figure 7C); this was named *BrFLC1as1022* based on its length. We used the same transcriptome data to analyze the expression level of *BrFLC1as1022*. It turned out that the expression of *BrFLC1as1022* was highest in NV and that the expression level continually declined during vernalization (Figure 7D).

**Figure 7.**
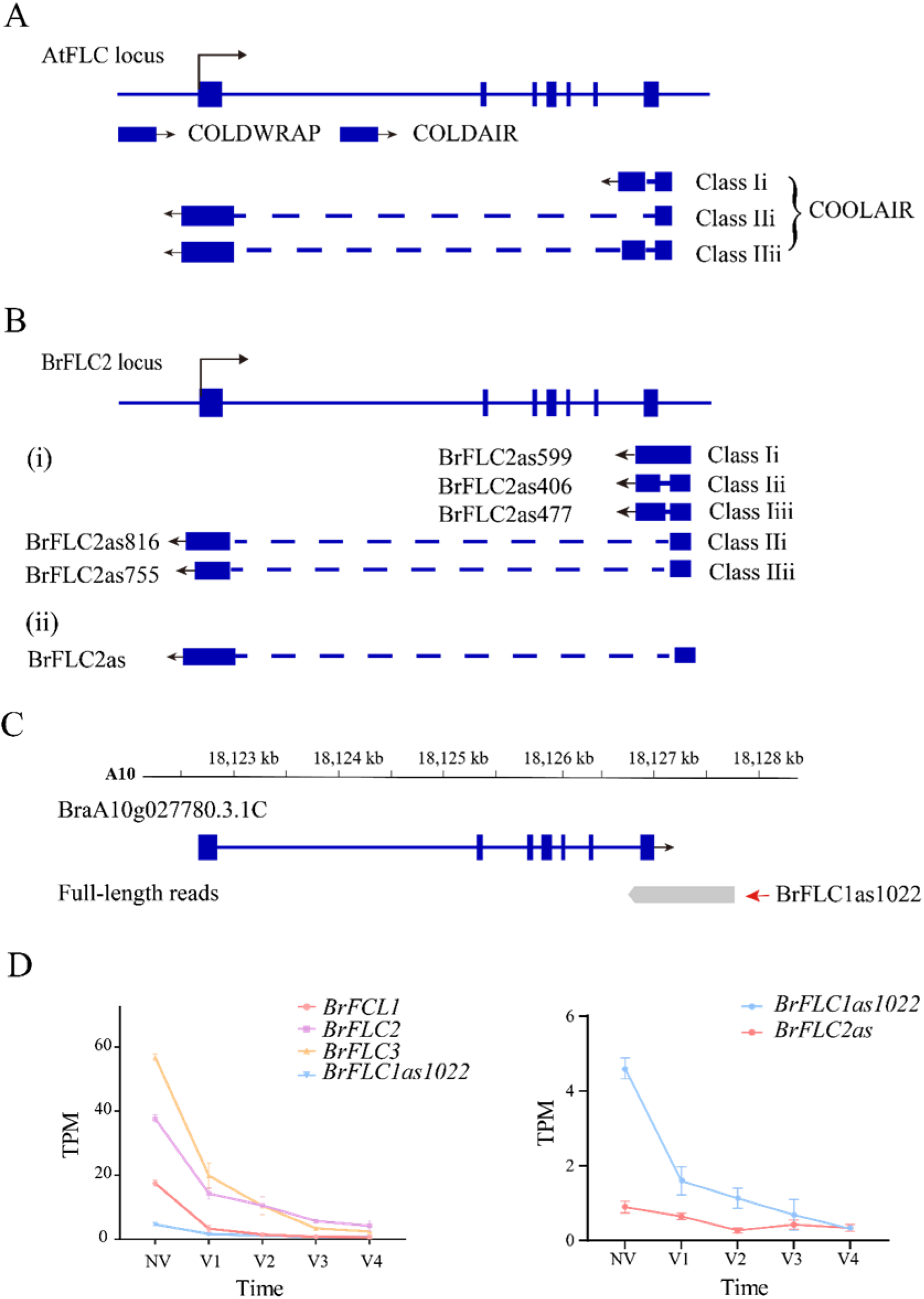
LncRNA identification in Iso-seq data. **(A)** *AtFLC* gene regulation by lncRNAs. *COOLAIR* is alternatively spliced. Class I and class II are abundant *COOLAIR* isoforms. The blue boxes indicate gene exons. The full line indicates chromosome. The dotted line indicates introns. The arrow mark indicates the direction of transcription. **(B)** *BrFLC2* gene regulation by *COOLAIR* in the previous studies. **(C)** The gene structure of *BrFLC1* in previous annotation and full-length transcript. The first of read is *PB.25365.1*. The second of read is *PB.25365.2* (*BrFLC1as1022*). **(D)** The line chart depicts the expression trend of *BrFLCs*, *BrFLC1as1022* and *BrFLC2as* during vernalization.

### Update of *B. rapa* reference genome annotation

In summary, in this study, we relied on the *B. rapa* genome to upgrade genome annotation based on the v3.1 annotation version, including refining the gene structure of 20,340 previously annotated genes, identifying gene UTR regions, and adding unannotated non-coding exons. We corrected 886 wrongly split genes into 427 genes.

A set of 830 novel genes missed in the previous annotation was eventually combined with 25,652 annotated genes that were not captured by full-length sequencing, resulting in a total of 47,249 coding genes in the new v3.5 annotation and released in BRAD (www.brassicadb.cn).

## DISCUSSION

Continuous refinement of annotation is a prerequisite for correctly interpreting the functional elements of the genome. Taking advantage of long read lengths from the PacBio sequencing technology, we generated a comprehensive full-length transcript dataset of *B. rapa* by pooling RNA from five tissues to construct two cDNA libraries. The new v3.5 annotation consists of 47,249 protein coding genes (830 novel genes), 13,550 TE containing loci, and 1,480 lncRNAs. The annotated loci span 118.6 Mb, an increase of 17.1 Mb from the previous *B. rapa* reference genome annotation v3.1.

The previous *B. rapa* annotations included only a single transcript for each gene, with no information concerning AS isoforms. In this study, we identified 28,564 AS events for 11,870 protein coding genes from Iso-Seq data without the aid of assembly. These genes accounted 25.1% of the total number of annotated genes in v3.5, providing valuable information for functional analysis and better explaining the richness of gene expression. The ratio of genes with AS isoforms was lower than that reported in Araport11 of *A. thaliana* (Cheng et al., 2017). The estimated frequency of AS forms increases along with the sampling depth, i.e., with the average number of transcripts sequenced for a gene (Zavolan et al., 2003). We believe that annotation of AS isoforms in this study is not saturated, since we only used quite limited types of tissue from plants grown only under normal or vernalization conditions for PacBio sequencing.

In this study, we found that the isoform number was positively correlated with both exon number and gene expression level. This differed from previous reports. There was a positive correlation between the number of exons and isoforms, but the mean expression levels of AS genes were slightly lower than those of non-AS genes in strawberry (Yuan et al., 2019). In maize, the number of isoforms in each gene was positively correlated with the number of exons but had no relationship with the gene expression level (Wang et al., 2016). However, we found that the number of isoforms per gene was positively correlated with exon number (*Pearson*, r = 0.295, *P*-value <0.001) as well as the expression level (*Pearson*, r = 0.124, *P*-value < 0.001) in *B. rapa*. We deduced that along with the increase of exon numbers, the number of splicing sites increases, leading to the formation of more isoforms. From the existing data, we found a total of 256 splicing motifs, among which the main type was the typical GT-AG pattern, accounting for 41.5%, while most of the other splice junctions accounted for less than 1% each. Moreover, such atypical splicing junctions generally occurred in genes with multiple exons (number > 5).

In previous studies, 2,237 (Paul et al., 2016) and 2,088 (Shea et al., 2019) lncRNAs were identified in *B. rapa* using cDNA sequences and Illumina short reads, respectively. In this study, we identified 1,480 lncRNAs in total, fewer than in previous reports. Since the sequences of the lncRNAs reported by Paul et al could not be accessed, we only compared 1,480 lncRNAs with the 2,088 lncRNAs reported by Shea et al (2019). The blast results showed similarities (*E-*value < 10e^−6^) in a total of 636 lncRNAs, including 559 lincRNAs, 66 NATs and 11 incRNAs. Shea et al (2019) compared expression level of lncRNAs in Chinese cabbage leaves with and without vernalization and detected 410 of the 1,444 lincRNAs, 120 of the 551 NATs, and 19 of the 93 incRNAs that was differentially expressed. In our study, we detected many fewer differentially expressed lncRNAs between vernalized and non-vernalized plants. However, the detection of the 209 lncRNAs showed a unique differential expression pattern in response to cold stress, suggesting that these lncRNAs are closely involved in governing the cold-stress responses of *B. rapa*.

*COOLAIR* has been suggested to play an important role in the regulation of *FLC* during vernalization in *A. thaliana* (Hawkes et al., 2016). Li et al (2016) isolated five NATs for *BrFLC2* by RACE-PCR and grouped them into two classes according to polyadenylation sites in intron 6 or within the promoter of *BrFLC2*. The NAT *BrFLC1as1022* we detected at the *BrFLC1* locus by Iso-Seq was polyadenylated within intron 6; therefore, it should belong to Class I. In the study (Shea et al., 2019), a COOLAIR-like transcript at the *BrFLC2* locus was detected and named as *BrFLC2as*. The expression of *BrFLC2as* was up-regulated upon short-term vernalization. We evaluated the expression of *BrFLC2as* using our RNA-seq data; the results showed that *BrFLC2as* expression decreased along with the extension of vernalization period similar to that of *BrFLC1as1022* detected by Iso-Seq (Figure 7D). Hawkes et al (2016) reported the structural conservation of *COOLAIR* across Brassicaceae species, and proposed a functional role for COOLAIR transcripts rather than, or in addition to, antisense transcription. To our knowledge, *BrFLC1as1022* is the first reported NAT for *BrFLC1.* This finding refutes the hypothesis proposed by Li et al (2016), that *BrFLC2* NATs contribute to the repression of all of the *BrFLC* genes in *B. rapa*.

Taken together, this work greatly improves existing gene models in *B. rapa*. Despite the substantial improvement in genome annotation v3.5, the work should be ongoing. With the aid of advances in sequencing technology and bioinformatics analysis, and additional samples from developmentally specific stages and different environmental stimuli, discovery of novel features will be included to update *B. rapa* genome annotation in the future.

## Supplementary Material

### Supplementary Table 1

Data information collected from public databases

### Supplementary Table 2

Gene-specific primers used for RT-PCR

### Supplementary Table 3

Statistics of the exon number of different gene types

### Supplementary Table 4

The percentages of splicing in each subgenome

### Supplementary Table 5

List of lincRNAs, NATs, and incRNAs

## Data Availability Statement

The Iso-Seq raw reads and Illumina sequencing data reported in this study were deposited in the NCBI Sequence Read Archive (SRA) under accession numbers SUB10710576 and SUB10725709.

## Author Contributions

W-XW and JW conceived the study. Z-CZ and CX performed data collection and bioinformatics analysis. JG and XX participated in Chiifu culture processing, RNA extraction and sequencing. L-YF conducted RT-PCR experiments. JW and Z-CZ wrote the manuscript. JW, X-WW, R-ML, J-LL and Z-CZ revised the manuscript. All authors read and approved the final manuscript.

## Funding

This work was funded by National Natural Science Foundation of China (32072593).

## Notes

### Competing Interest Statement

The authors have declared no competing interest.

